# UNC-10/SYD-2 links kinesin-3 to RAB-3-containing vesicles in the absence of the motor’s PH domain

**DOI:** 10.1101/723247

**Authors:** Odvogmed Bayansan, Prerana Bhan, Chien-Yu Chang, Syed Nooruzuha Barmaver, Che-Piao Shen, Oliver Ingvar Wagner

## Abstract

Kinesin-3 KIF1A (UNC-104 in *C. elegans*) is the major axonal transporter of synaptic vesicles and mutations in this molecular motor are linked to KIF1A-associated neurological disorders (KAND) including Charcot-Marie-Tooth disease, amyotrophic lateral sclerosis and hereditary spastic paraplegia. UNC-104 binds via its PH (pleckstrin homology) domain to the lipid bilayers of membranous vesicles which is considered a weak interaction. RT-PCR and Western blot experiments reveal genetic relations between SYD-2, UNC-10 and RAB-3. Co-immunoprecipitation assays reveal functional relations and bimolecular fluorescence complementation (BiFC) assays expose *in situ* interactions between these proteins. Though both SNB-1 and RAB-3 are actively transported by UNC-104, the movement of RAB-3 is generally enhanced and largely depending on the presence of SYD-2/UNC-10. Deletion of UNC-104’s PH domain did not affect UNC-104/RAB-3 colocalization but did affect UNC-104/SNB-1 colocalization. Similarly, motility of RAB-3-labeled vesicles is unaltered in nematodes carrying a point mutation in the PH domain while movement of SNB-1 is significantly reduced in anterograde directions. These findings suggest a dual UNC-10/SYD-2 linker acting as a sufficient buttress to connect the motor to RAB-3-containing vesicles to enhance their transport. This additional linker will also strengthen the rather weak motor-lipid interaction.

**Graphical abstract:** 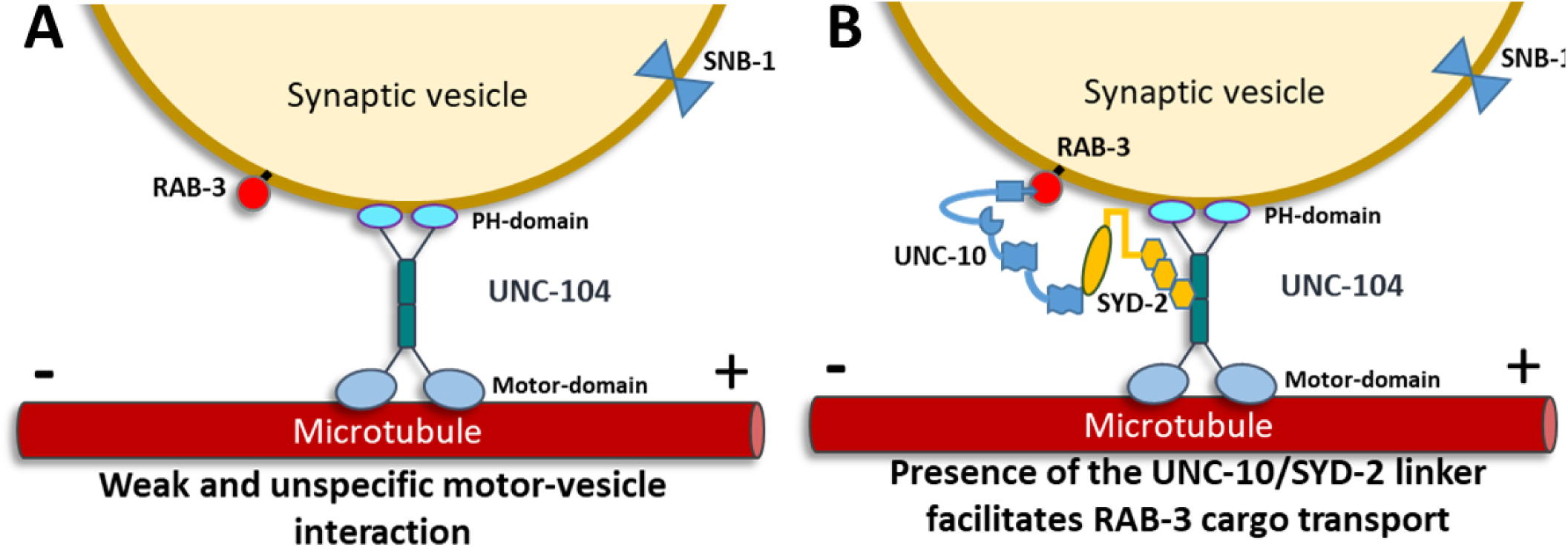

## INTRODUCTION

Understanding the molecular basis of axonal transport is critical to develop novel drugs to prevent or cure various neurological diseases such as Charcot–Marie–Tooth disease (CMT), Amyotrophic lateral sclerosis (ALS) or Alzheimer’s disease (Guillaud et al., 2020; Guo et al., 2020; Surana et al., 2020). The neuronal cytoskeleton is composed of microtubules (MTs), actin filaments as well as neurofilaments (NFs) all functioning to maintain neuronal polarity, morphology and integrity of axons (Barlan and Gelfand, 2017; Brady and Morfini, 2017). Axonal transport supplies axons and nerve terminals with proteins, lipids and mitochondria and prevents the build-up of misfolded proteins and toxic aggregates (Guedes-Dias and Holzbaur, 2019; Perlson et al., 2010). Kinesins move from the minus-ends to plus-ends of uniformly oriented MTs in axons (anterograde transport) thereby transporting STVs (synaptic vesicle protein transport vesicles) from neuronal cell bodies (somas) to axonal termini. Retrograde transport (from the minus-ends to the plus-ends of MTs) is solely accomplished by cytoplasmic dynein (Fan and Lai, 2022; Guedes-Dias and Holzbaur, 2019). KIF1A is a neuron-specific molecular motor belonging to the kinesin-3 family and is termed as UNC-104 in the model organism *C. elegans.* Mutations in kinesin-3 are associated to various neurological disorders including CMT, ALS and hereditary spastic paraplegia (HSP) (Ali and Yang, 2020; Brady and Morfini, 2017; Millecamps and Julien, 2013). KIF-1A/UNC-104 is the major axonal transporter of synaptic precursors such as RAB-3, synaptotagmin, synaptobrevin and synaptophysin (Hall and Hedgecock, 1991; Hayashi et al., 2019; Maeder et al., 2014). UNC-104 has been shown to transport dense core vesicles (Zahn et al., 2004) and to regulate size and density of synapses (Niwa et al., 2016). Besides its function to regulate dendrite development (Kern et al., 2013; Zhang et al., 2016), KIF1A/UNC-104 also plays a role in synaptic aging and memory (Li et al., 2016). Recently, it has been shown that KIF-1A is required for localization of the presynaptic adhesion molecule neurexin in GABAergic dendritic spine formation and synapse maturation (Oliver et al., 2022). Structurally, KIF-1A/UNC-104 is composed of six major domains, including the motor head, fork head associated (FHA), pleckstrin homology (PH), coiled coils 1, 2 and 3 domains (CC) (Hirokawa et al., 2009). In *C. elegans*, UNC-104 associates with tau (termed PTL-1 in worms) and mutations in *ptl-1* facilitate SNB-1 (synaptobrevin-1) cargo movement (Tien et al., 2011). If not bound to cargo, kinesin-3 motors appear either monomeric or as inactivated dimers via intramolecular folding. Upon cargo-binding, kinesin-3 motors rapidly dimerize to become processive for fast and directed transport (Akhmanova and Hammer, 2010; Hammond et al., 2009; Hummel and Hoogenraad, 2021; Rashid et al., 2005). Kinesin-3 monomeric motors also quickly dimerize in lipid rafts (found in vesicular surfaces) due to their ability to bind to phosphatidylinositol 4,5-bisphosphate (PtdIns(4,5)P_2_) via their PH domains (Klopfenstein and Vale, 2004).

Active zones are the sites for vesicle docking and fusion at synapses involving proteins MUNC-13 (UNC-13), RAB3A (RAB-3), RIMS1 (UNC-10), liprin-α (SYD-2), CASK (LIN-2) or Clarinet (CLA-1) (Krout et al., 2023). Direct and functional interactions between these active zone components and UNC-104 have been reported such as UNC-104/SYD-2 (Wagner et al., 2009), UNC-104/RAB-3 (Nonet et al., 1997), UNC-104/CLA-1 (Xuan et al., 2017), UNC-104/NRX-1 (Oliver et al., 2022) and UNC-104/LIN-2 (Wu et al., 2016). SYD-2 is a multi-domain scaffolding protein that encompasses 8 N-terminal coiled coils and 3 C-terminal sterile alpha motifs (SAM domains) which are able to interact with each other to regulate protein function via intramolecular folding (Chia et al., 2013). In *C. elegans syd-2* mutants, active zones appear diffuse and more irregular along axons causing impairment of synaptic vesicle docking, thus, synaptic transmission (Kittelmann et al., 2013; Stigloher et al., 2011; Zhen and Jin, 1999). The N-terminal domain of SYD-2 directly associates with other synaptic proteins such as CAST/ELKS, GIT1 and RIMS1(UNC-10) (Ackermann et al., 2015; Dai et al., 2006; Ko et al., 2003; Shin et al., 2003; Spangler et al., 2013), while the C-terminal SAM domains interact with the FHA domain of UNC-104 to activate the motor (Wagner et al., 2009). SYD-2/RAB-3 interactions are reported to maintain the synaptic vesicle pool at presynaptic dense projections (Wu et al., 2013).

RIMs are a family of multi-domain scaffolding proteins that were initially identified as sole Rab3-effectors to regulate synaptic vesicle fusion (Wang et al., 1997). The RIMS1 homolog in *C. elegans* is UNC-10 (Koushika et al., 2001) composed of an N-terminal zinc finger motif, a central PDZ domain, two C-terminal C2 domains (C2A and C2B) and a proline-rich SH3 motif (Wang et al., 2000). RIMS1/UNC-10 interacts with multiple synaptic proteins, e.g., its N-terminal zinc finger domain interacts with GTP-bound Rab3 (Wu et al., 2023), while its C-terminal C_2_B domain associates with the N-terminus of Liprin-α/SYD-2 (Schoch et al., 2002). Liprin-α regulates the presynaptic organization by anchoring RIM1 and CASK to facilitate synaptic vesicle release (Spangler et al., 2013) and it has been shown that UNC-10 and SYD-2 act in the same pathway to regulate synaptic vesicle turnover (Stigloher et al., 2011). Besides, UNC-10/RIM and SYD-2/liprin-α are critical for presynaptic localization of voltage-gated calcium channels to the exocytosis machinery at the presynaptic active zones in *C. elegans* (Oh et al., 2021).

Rab3 is a member of the RAB family of small GTPases that recruits synaptic vesicles for vesicle docking and fusion at active zones (Geppert et al., 1994). It integrates into the synaptic vesicle membrane via its C-terminal sequence Cys-X-Cys that is highly conserved in invertebrates (Johnston et al., 1991). Rab3A binds to synaptic vesicles in its GTP-bound state and dissociation occurs after GTP-hydrolysis (Fischer von Mollard et al., 1994). The guanine nucleotide exchange factor (GEF) AEX-3 (DENN/MADD) plays an important role in regulating RAB-3, concomitantly affecting proper localization of presynaptic proteins and neurotransmitter release (Mahoney et al., 2006). Rab3A is transported to the active zone by the fast anterograde axonal transport system (Li et al., 1995) and in mammals KIF1A associates to Rab3-carrying vesicles via DENN/MADD (Niwa et al., 2008). Rab3A null mutant mice (Geppert et al., 1994) and worms (Nonet et al., 1997) display mild behavioral abnormalities, with worms specifically displaying resistance to the pesticide aldicarb. Also, synaptic transmission is perturbed in *C. elegans rab-3* mutants with a reduced number of synaptic vesicles near the active zones (based on transport defects) (Nonet et al., 1997).

Due to the characterized weak and non-specific binding between PH domains and lipid bilayers (Lemmon and Ferguson, 2001), and based on the aforementioned knowledge from the literature, we propose that UNC-10/SYD-2 would act as a novel receptor for UNC-104-mediated transport of RAB-3 containing vesicles. At the same time, the RAB-3/UNC-10/SYD-2 linkage would reinforce the weak motor-lipid interaction.

## RESULTS

### Genetic relation between *syd-2*, *unc-10* and *rab-3* in nematodes

To understand the genetic relation between SYD-2, UNC-10 and RAB-3 in nematodes, we investigated RNA and protein expression changes of SYD-2 in various *unc-10* and *rab-3* genetic backgrounds. Interestingly, syd-2 expression levels significantly increased in *C. elegans* carrying mutations in the unc-10 gene but decreased in a *rab-3* mutant (Fig. 1A+B). The strain NM1657 *unc-10*(*md1117*) carries a large 8601 bp deletion in the unc-10 gene that includes the entire coding region along with regions of the 5’ and 3’ UTRs, thus likely being a knockout (Suppl. Fig. S1B, (Koushika et al., 2001)). Strain CB102 *unc-10*(*e102*) carries a C to T substitution in intron 15 of the *unc-10* gene resulting in splicing defects that produce a truncated protein excluding the C-terminal domain (Suppl. Fig. S1B, (Koushika et al., 2001)). The phenotypes of both strains NM1657 and CB102 are “mild coiler” and “mild uncoordinated”. The strain NM791 *rab-3*(*js49*) carries a G to A transition at position 2 in the tryptophan 76 codon in the *rab-3* gene leading to a nonsense mutation causing a premature stop likely being a knockout (Suppl. Fig. S1B, (Nonet et al., 1997)). The worm phenotype of NM791 is “strong coiler”. Note that two other strains NM210 and NM211 exist carrying *rab-3* alone, however, these strains are poorly characterized and worm phenotypes are less pronounced (only mild uncoordinated) (wormbase.org), thus we refrained from employing them for this study. As another result, protein expression goes well along with changes in RNA levels: with increasing RNA expression of SYD-2 in *unc-10* mutants (Fig. 1A), protein expression of SYD-2 increased as well (Fig. 1C); and with decreasing RNA levels in of SYD-2 in *rab-3* mutants, expression levels of SYD decreased as well (Fig. 1D).

**Figure 1:**
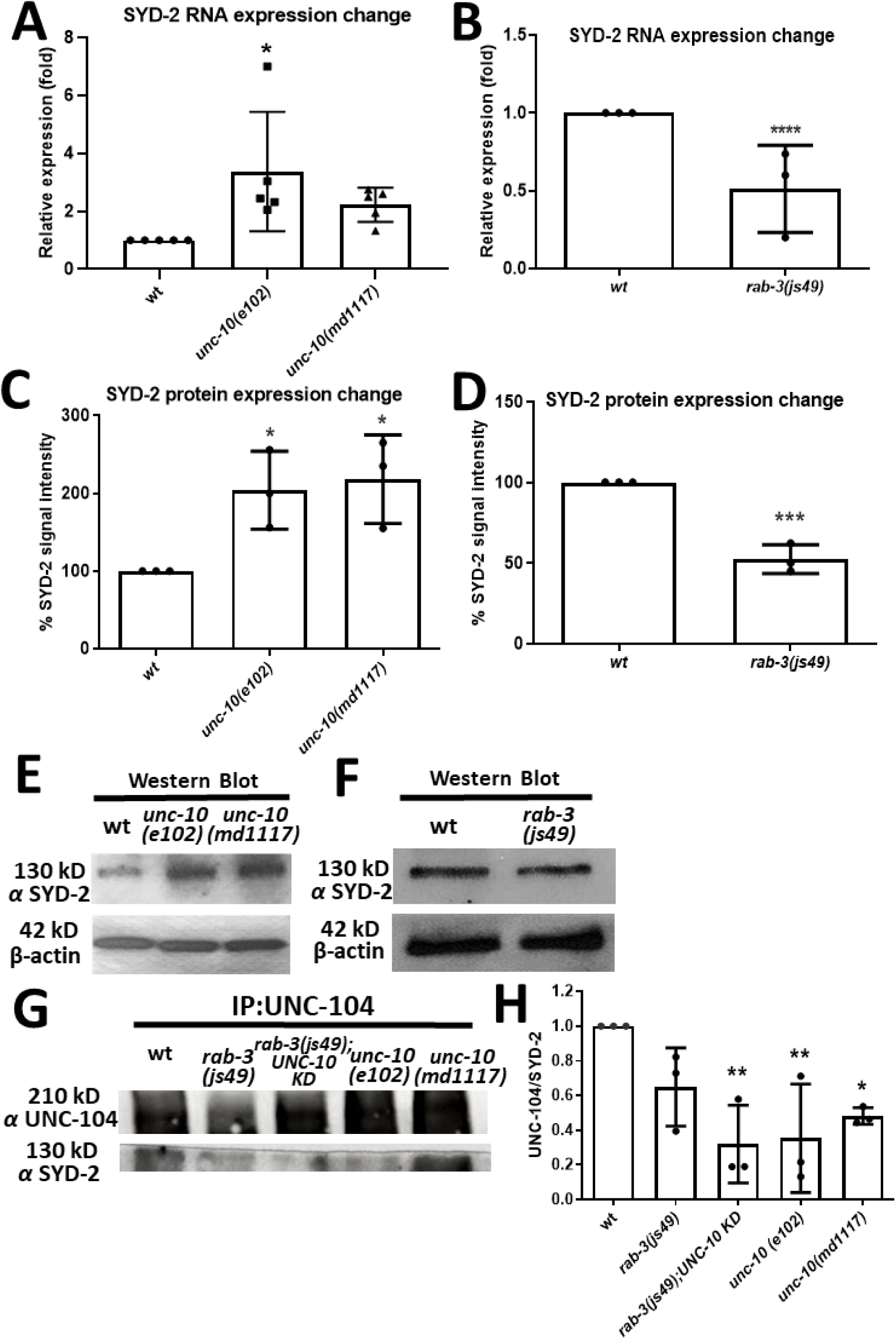
Genetic and functional relations between *syd-2, unc-10* and *rab-3* in *C. elegans* animals. (A+B) Relative syd-2 RNA expression levels in (A) *unc-10* and (B) *rab-3* allelic backgrounds. RNA levels normalized to endogenous cdc-42 expression as an internal control. (C+D) Quantification of data from Western blots shown in (E+F). (G) Co-immunoprecipitation assay employing lysates from whole worms expressing UNC-104::mRFP in wt, *unc-10*, *rab-3* and *rab-3*;*unc-10 KD* backgrounds. (H) Quantification of Co-IP shown in (G). (A), (C) and (H): One-way ANOVA with Dunnett’s test for multiple comparisons with *p<0.05, **p<0.005, ***p<0.001. (B+D): Two-tailed, unpaired Student’s t-test with ***p<0.001. Graphs represent mean ± STD. Number of trials for each experimental group = 3.

### Effects of *unc-10* and *rab-3* mutations on the UNC-104/SYD-2 complex

The functional interactions between SYD-2 and UNC-104 have been well described in the literature as summarized above. To understand the role of UNC-10 and RAB-3 in this interaction scheme, we carried out co-immunoprecipitation assays. Precipitating UNC-104 using an anti-UNC-104 antibody from whole worm lysates, we found that bound SYD-2 protein content was reduced in both *unc-10*(*md1117*) and *unc-10*(*e102*) mutants (Fig. 1G+H). Though SYD-2 expression remained unchanged in *rab-3*(*js49*) single mutants, expression of SYD-2 was significantly reduced in *rab-3*(*js49*);*unc-10 KD* double mutants (Fig. 1G+H). These data point to a functional relation between the UNC-104/SYD-2 complex and UNC-10 as well as RAB-3. To understand whether UNC-104/SYD-2 interactions also change in neurons of *unc-10* or *rab-3* worms, we performed colocalization as well as bimolecular fluorescence complementation (BiFC) assays. BiFC is a well-established assay in living *C. elegans* animals for investigating *in situ* functional relations between two proteins. Here, the YFP protein Venus is split into two halves (N-terminal Venus, VN, and C-terminal Venus, VC) and the test proteins are fused to each half. Complementation of Venus, leading to yellow fluorescent signals, occurs only if the two test proteins are at least 7-10 nm close to each other, therefore likely forming functional complexes (Hsu et al., 2011; Hu and Kerppola, 2003). Our data reveal that colocalization between UNC-104 and SYD-2 in ALM neurons (Suppl. Fig. S1A) significantly decreases in *unc-10*(*md1117*) and *rab-3*(*js49*) mutants as well as in *rab-3*(*js49*);*unc-10 KD* double mutants (Fig. 2A+B, Suppl. Fig. S2 for nerve ring). More importantly, physical interactions between UNC-104/SYD-2 complexes (as revealed by BiFC experiments) are disrupted in *unc-10*(*md1117*) and *rab-3*(*js49*) mutants (Fig. 2C+D). These *in situ* data are consistent with our *in vitro* observations from Figure 1 and support our hypothesis that UNC-104/SYD-2 complexes are dependent on the presence of UNC-10 and RAB-3.

**Figure 2:**
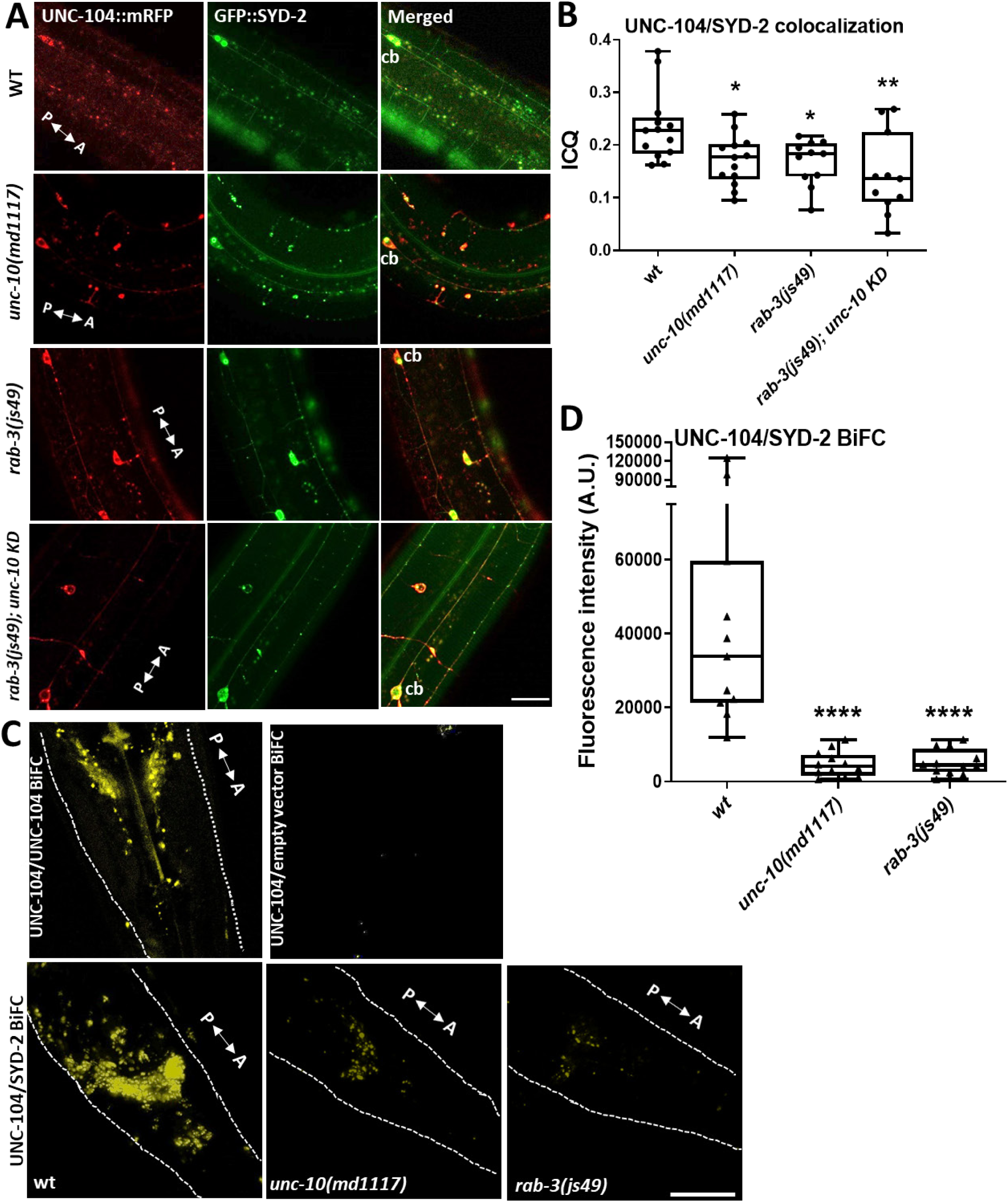
Colocalization and bimolecular fluorescence complementation assays (BiFC) between UNC-104 and SYD-2 in the presence or absence of *unc-10* or *rab-3* mutations. (A) UNC-104 and SYD-2 colocalization in ALM neurons (Suppl. Fig. S1A) of wild type, *unc-10*, *rab-3*, and *rab-3;unc-10 KD* worms. (B) Intensity correlation coefficients from images shown in (A) (n = 12 worms). (C) UNC-104 and SYD-2 BiFC signals in the nerve ring of wild type, *unc-10* and *rab-3* worms. UNC-104/UNC-104 BiFC used as positive and UNC-104/empty (empty vector) used as negative BiFC control. White dashed lines designate the borders of the worm’s head (Suppl. Fig. S1A). (D) Quantification of BiFC signals shown in (C) (n = 12 worms). A = anterior, P = posterior directions. Cb = cell body of ALM neuron. One-way ANOVA with Dunnett’s test for multiple comparisons with *p<0.05, **p<0.005, ***p< 0.001, ****p<0.0001. Box and whisker graphs represent maximum value, upper quartile, median, lower quartile and minimum value. Scale bars: 10 µm.

### Role of *syd-2*, *unc-10* and *rab-3* single/double mutations on UNC-104 particle distribution in neurons

To understand the physiological consequences of the tight interplay between these proteins, we measured changes in neuronal UNC-104 particle distribution in single (*syd-2*, *unc-10* and *rab-3*) and in double mutants (*syd-2*/*unc-10*, *syd-2*/*rab-3*, *rab-3*/*unc-10*). In a previous study, we reported that UNC-104 forms SYD-2-dependent cluster along axons in an arrangement that does not match known *en passant* synapse patterns, and we also demonstrated that such clusters are dynamic structures rather than inactive motor aggregates (Wagner et al., 2009). Here, we evaluate UNC-104 cluster formation in the sublateral system as well as in the long mechanosensory neuron ALM (Suppl. Fig. S1A). Though the focus lies in analyzing the single ALM neuron, it is evident that in both types of neurons (sublaterals and ALM) UNC-104 cluster become significantly smaller in *unc-10* and *syd-2* mutants (as well as in *rab-3* mutants when inspecting ALM alone) (Fig. 3A-C and E; Suppl. Table S1+2). Critically, these reductions in UNC-104 cluster sizes can be reversed to wild type levels via rescue experiments in which wild type genes are expressed in the respective mutants (Fig. 3E, Suppl. Table S2). Though double mutants did not induce clusters of even smaller sizes (as opposed to single mutants) they, on the other hand, also neither rescued nor enhanced the effects as observed in single mutants (except for *syd-2*;*unc-10* double mutant) (Fig. 3C+E). We also analyzed how far UNC-104 particles travel from the axon hillock to distal regions of the ALM neuron (Suppl. Fig. S1C). From Figure 3C+D (Suppl. Table S2), it is obvious that travel distances are significantly reduced in *syd-2* mutants and that this effect can be rescued by overexpressing the wild type gene in the mutants. While such an effect cannot be seen in single *rab-3* mutants, in *rab-3*;*unc-10* as well as in *syd-2;unc-10* double mutants travel distances are significantly reduced. Interestingly, in *rab-3*;*syd-2* mutants travel distances even increase pointing to an unknown compensatory pathway. Nevertheless, that nearly all double mutants reveal significant different effects (as compared to the single mutants) indicates the strong interdependences of these proteins in regulating UNC-104 particle size and distribution. Especially that *rab-3* further decreases UNC-104 travel distances in *unc-10* mutants is in accordance with our model of RAB-3 being tightly linked to UNC-10. Because we assume that the reduction of UNC-104 travel distances is related to diminished motor speeds, we therefore analyzed UNC-104 motility parameters.

**Figure 3:**
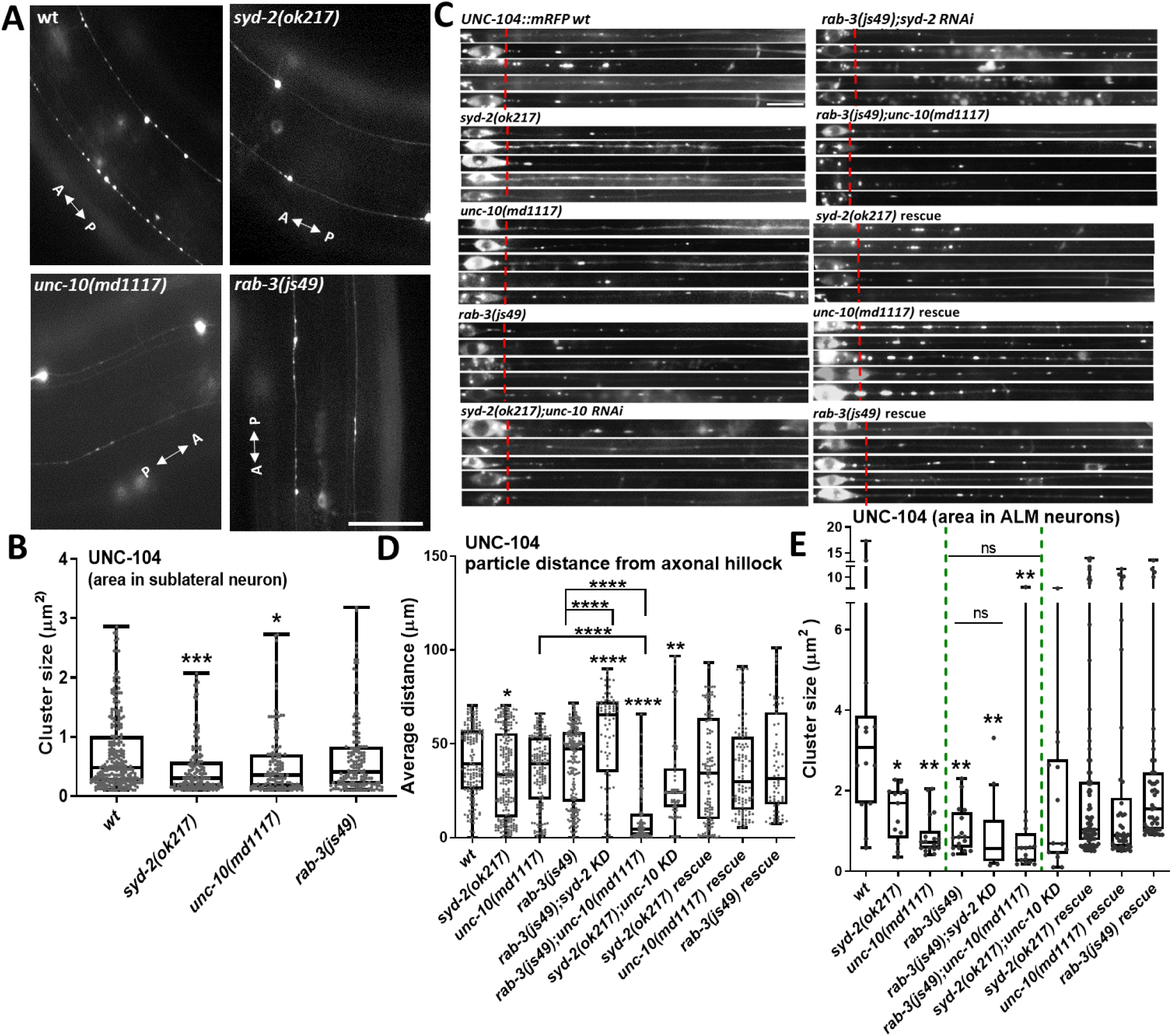
Effect of *syd-2*, *unc-10* and *rab-3* single and double mutants on UNC-104 clustering and travel distances in sublateral and ALM neurons. (A) UNC-104::mRFP cluster patterns in sublaterals nerve bundles (Suppl. Fig. S1A). (B) Cluster size (area) quantification from (A). (C) Stacks of ALM neurons (after digital straightening) to examine UNC-104 cluster patterns in different mutants. (D) Quantification of UNC-104 particles travelled from the axonal hillock (red dotted line in (C)) to distal areas of the neuron (data taken from (C)) (Suppl. Fig. S1C). (E) Cluster size (area) quantification from (C). One-way ANOVA with Dunnett’s test for multiple comparisons with *p<0.05, **p<0.005, ***p< 0.001, ****p<0.0001. For two group comparisons in (D) two-tailed, unpaired Student’s t-test with ****p<0.0001 was used. Box and whisker graphs represent maximum value, upper quartile, median, lower quartile and minimum value. N = 20 worms per experimental group. Further details of data can be found in Suppl. Tables S1+2. Scale bar: 10 µm.

### The role of *syd-2*, *unc-10* and *rab-3* single/double mutations on UNC-104 motility

To monitor UNC-104 motility in the long (up to 500 μm) ALM neurons, we tagged the motor’s C-terminal end with mRFP that is reported to be less prone to self-aggregation as opposed to GFP (Campbell et al., 2002). From kymograph analysis, we calculate various motility parameters such as velocities, run lengths (event run lengths and total run lengths) as well as pausing (Suppl. Fig. S1D). As a result, all single genes (allelic mutations or gene knockdown) *syd-2*, *unc-10* and *rab-3* significantly reduce UNC-104’s moving speeds in anterograde directions (Fig. 4A, Suppl. Fig. S3A+B, Suppl. Table S3). Similarly, knocking down *syd-2* and *unc-10* resulted in reduced UNC-104 velocities. Intriguingly, UNC-104’s speed is further decreased in double mutants *rab-3(js49);syd-2 KD* and *rab-3(js49);unc-10 KD* compared to single mutant *rab-3(js49)*. Interestingly, anterograde event run lengths and total run lengths are significantly decreased in *unc-10(md1117)* knockout allele but not in *unc-10(e102)* point mutation allele. Also, *rab-3* and *syd-2 KD* (but not *ok217* allele) both negatively affect UNC-104 run lengths (Fig. 4 B+C, Suppl. Fig. S3A+C, Suppl. Table S3). Strikingly, SYD-2 overexpression led to increased UNC-104 run lengths even in *unc-10 KD* backgrounds (Fig. 4B+C) consistent with the literature reporting SYD-2 as an UNC-104 activator (Muniesh et al., 2020; Wagner et al., 2009; Wu et al., 2016). Furthermore, velocities as well as motor run lengths were significantly reduced in *rab-3* double mutants *rab-3*;*unc-10 KD* and *rab-3*;*syd-2 KD*, once more pointing to functional, synergistic effects between these genes. Notably, reduced velocities and reduced run lengths in the *rab-3*;*unc-10 KD* double mutant are consistent with observations from Figure 3C+D revealing reduced travel distances for this double mutant. Reduced velocities concomitantly result in increasing pausing of the motor as measured in *unc-10(md1117), rab-3(js49)* and *rab-3(js49);unc-10 KD* mutants (Fig. 4D). Critically, we were able to successfully rescue all observed effects by overexpressing the respective wild type gene in the mutants (Fig. 4A-D). In conclusion, single and double mutant analysis point to strong synergistic effects between RAB-3, UNC-10 and SYD-2 to regulate UNC-104 motility.

**Figure 4:**
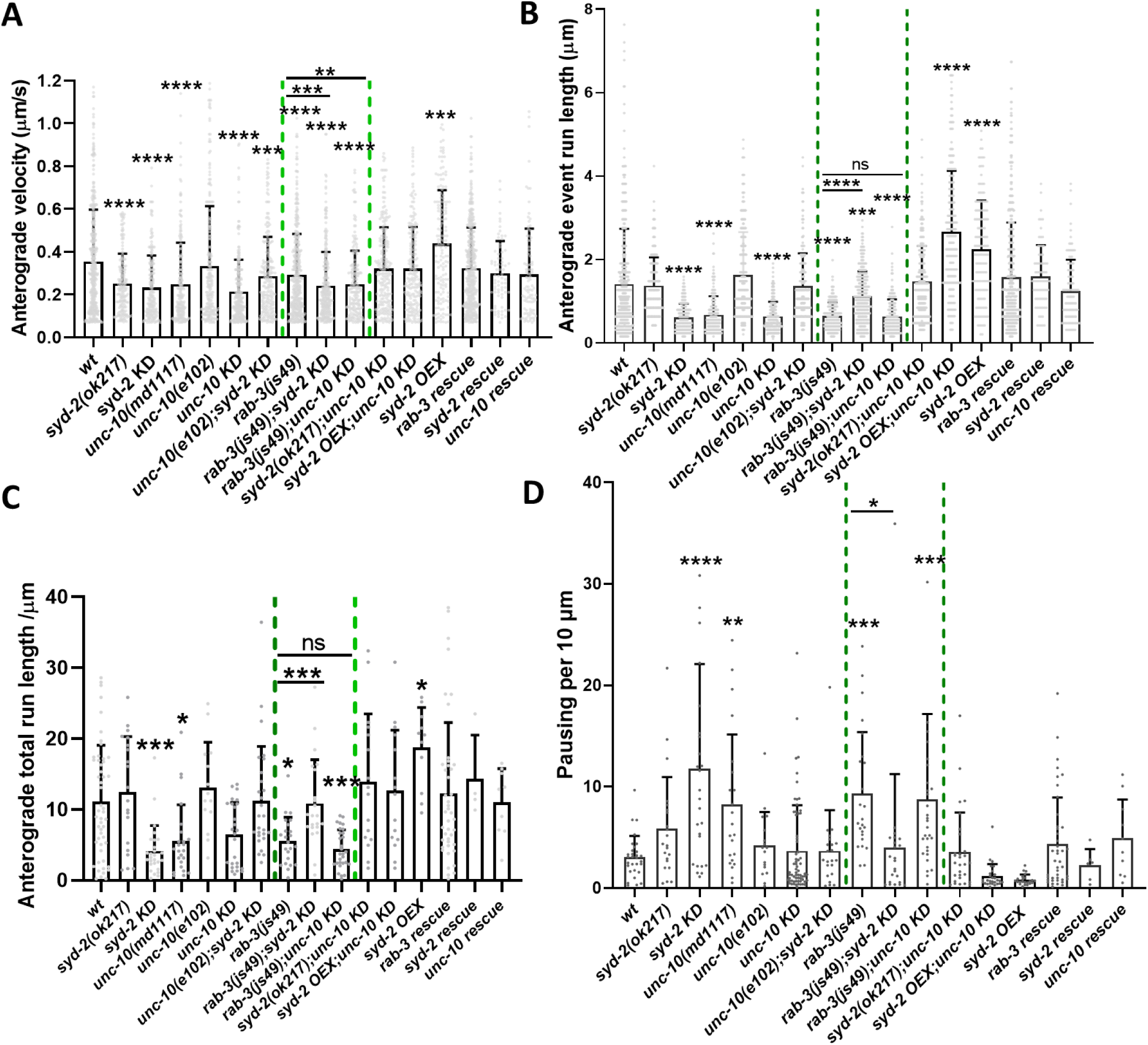
Effect of *syd-2*, *unc-10* and *rab-3* single and double mutants on UNC-104 motility in neurons of living *C. elegans* animals. (A) Anterograde velocity of UNC-104 in *syd-2, unc-10, rab-3* single mutants as well as in various double mutant combinations and rescues. (B) UNC-104 anterograde event run lengths. (C) UNC-104 anterograde total run length. (D) UNC-104 pausing per 10 µm. Green dotted lines outline particular data set comparisons. KD = knockdown, OEX = overexpression. Analyzed total events: UNC-104(wt) = 1531, *syd-2*(*ok217*) = 529, *syd-2 KD* = 697, *unc-10*(*md1117*) = 643, *unc-10 KD* = 740, *unc-10(e102);syd-2 KD* = 523, *rab-3*(*js49*) = 622, *rab-3;unc-10 KD* = 805, *syd-2*(*ok217*)*;unc-10 KD* = 536, r*ab-3*(*js49*)*;syd-2 KD* = 735, *rab-3* rescue = 1159, *syd-2* rescue = 155, *unc-10* rescue = 207, *syd-2 OEX* = 595, *syd-2 OEX;unc-10 KD* = 714. One-way ANOVA with Dunnett’s test for multiple comparisons with *p<0.05, **p<0.005, ***p< 0.001, ****p<0.0001. Two-tailed, unpaired Student’s t-test for comparisons of means of two groups with *p<0.05, **p<0.01, ***p<0.001, ****p<0.0001. Graphs represent mean ± STD. Further details of data can be found in Suppl. Fig. S3 and Suppl. Table S3.

### UNC-10/SYD-2 complex is required for RAB-3-but not for SNB-1-based transport

Based on above results, we hypothesize that the UNC-10/SYD-2 complex facilitate binding between UNC-104 and RAB-3-containing vesicles (Fig. 7). In this model, however, the UNC-10/SYD-2 dual linker complex would not affect binding between motor and SNB-1-containing vesicles. To approach this hypothesis, we analyzed motility as well as travel distances of both mCherry::RAB-3 and SNB-1::mRFP tagged synaptic vesicles in different mutant backgrounds (Fig. 5). The UNC-104-activating effect of SYD-2 on synaptic vesicle transport (both RAB-3- and SNB-1-bound vesicles) has been well reported (Muniesh et al., 2020; Wagner et al., 2009; Wu et al., 2016) and it is not surprising that in *syd-2*(*ok217*) knockout worms speeds of both SNB-1- and RAB-3-tagged vesicles are significantly reduced (Fig. 5A+C, Suppl. Table S4). Notably, speeds and travel distances of RAB-3-tagged vesicles are significantly decreased in double mutants *unc-10(md1117);syd-2 KD* while speeds and travel distances of SNB-1-tagged vesicles remain unaffected (Fig. 5, Suppl. Table S4+5). Interestingly, RAB-3-bound vesicles cover generally larger travel distances as compared to SNB-1-bound vesicles (Fig. 5B+D and E+F, wild type data). As a negative control, we knocked down *rab-3* in SNB-1::mRFP as well as *snb-1* in mCherry::RAB-3 expressing worms which, as expected, did not affect vesicle speeds (Fig. 5A+C). To demonstrate that the transport of RAB-3-containing vesicles is independent of UNC-104’s PH domain, we monitored RAB-3- and SNB-1-tagged synaptic vesicles in *unc-104(e1265)* mutants carrying a point mutation (D1497N) in the PI(4,5)P_2_ binding pocket of the PH domain. As expected, anterograde speeds were significantly decreased in SNB-1-tagged vesicles while speeds of RAB-3-tagged vesicles remain unchanged (Fig. 5A+C). In terms of anterograde travel distances, the point mutation in the PH domain of UNC-104 led to a 67% reduction in average distances (compared to wildtype) for SNB-1 movements but reduced to only 19% for RAB-3 movements (Fig. 5 B+D). These results are consistent with our model (Fig. 7) that transport of SNB-1 is dependent on UNC-104’s PH domain while the transport of RAB-3 depends only to a minor extent of this domain and is additionally stabilized by the UNC-10/SYD-2 linker.

**Figure 5:**
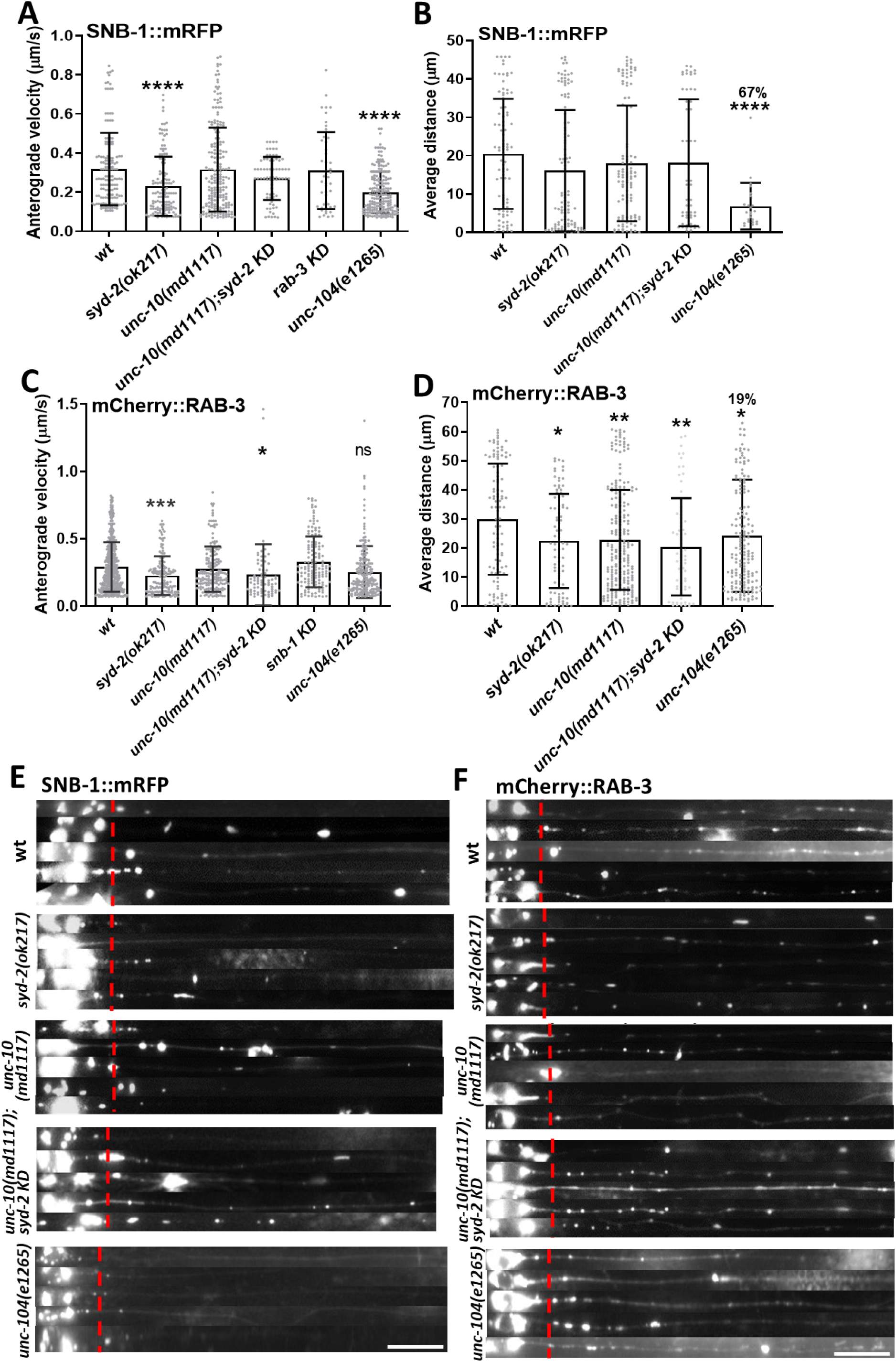
Effect of *syd-2*, *unc-10* and *rab-3* single and double mutants on SNB-1 and RAB-3 cargo trafficking. (A) Anterograde velocities of SNB-1-tagged synaptic vesicles. (B) Quantification of average distances travelled from the axonal hillock (red dotted line in (E)) to distal areas in the neuron (n = 20 worms). (C) Anterograde velocities of RAB-3-tagged synaptic vesicles. (D) Quantification of average distances travelled from the axonal hillock (red dotted line in (F)) to distal areas in the neuron. (E+F) Stacks of ALM neurons reveals characteristic SNB-1 (E) and RAB-3 (F) cluster patterns in different mutants. Further details of these data can be found in Suppl. Tables S4+5. Analyzed events in (A+B): SNB-1(wt) = 510, *rab-3 KD* = 360, *syd-2*(*ok217*) = 439, *unc-10*(*md1117*) = 518, *unc-10*(*md1117*)*;syd-2 KD* = 250, *unc-104(e1265) =* 628. Analyzed events in (C+D): RAB-3(wt) = 513, *snb-1 KD* = 306, *syd-2*(*ok217*) = 508, *unc-10*(*md1117*) = 509 and *unc-10*(*md1117*)*;syd-2 KD* = 300, *unc-104(e1265)* = 767. One-way ANOVA with Dunnett’s test for multiple comparisons with *p<0.05, **p<0.005, ***p< 0.001, **** p<0.0001. Graphs represent mean ± STD. Scale bar: 10 µm.

### Dual linker UNC-10/SYD-2 is sufficient to associate UNC-104 to RAB-3 bound vesicles in the absence of the motor’s PH domain

Based on our model, if eliminating UNC-104’s PH domain, the motor may still effectively transport RAB-3-bound vesicles. To adequately understand whether this assumption is valid, we designed worms double expressing UNC-104:GFP with a deleted PH domain as well as mCherry::RAB-3 or SNB-1::mRFP. As a result, colocalization between UNC-104 and RAB-3 (in sublateral neurons) remains unaffected in the absence of UNC-104’s PH domain (Fig. 6A+B and D, Suppl. Fig. S4A+B line scans) while colocalization between UNC-104 and SNB-1 is significantly reduced in the absence of the PH domain (Fig. 6E+F, Suppl. Fig. S4E+F line scans). This effect is equally evident in dorsal (Fig. 6G) and ventral (Fig. 6H) nerve cords (note that cell bodies of motor neurons are present in ventral nerve cords but not in dorsal nerve cords). While deleting UNC-104’s PH domain significantly reduces UNC-104/SNB-1 colocalization, *unc-10* knockout did not reduce UNC-104/SNB-1 colocalization (Fig. 6E+F, Suppl. Fig. S4G). On the other hand, colocalization between UNC-104 and RAB-3 is significantly reduced in *syd-2* and *unc-10* mutants (Fig. 6C+D, Suppl. Fig. S4C+D). These data support our model as summarized in Figure 7.

**Figure 6:**
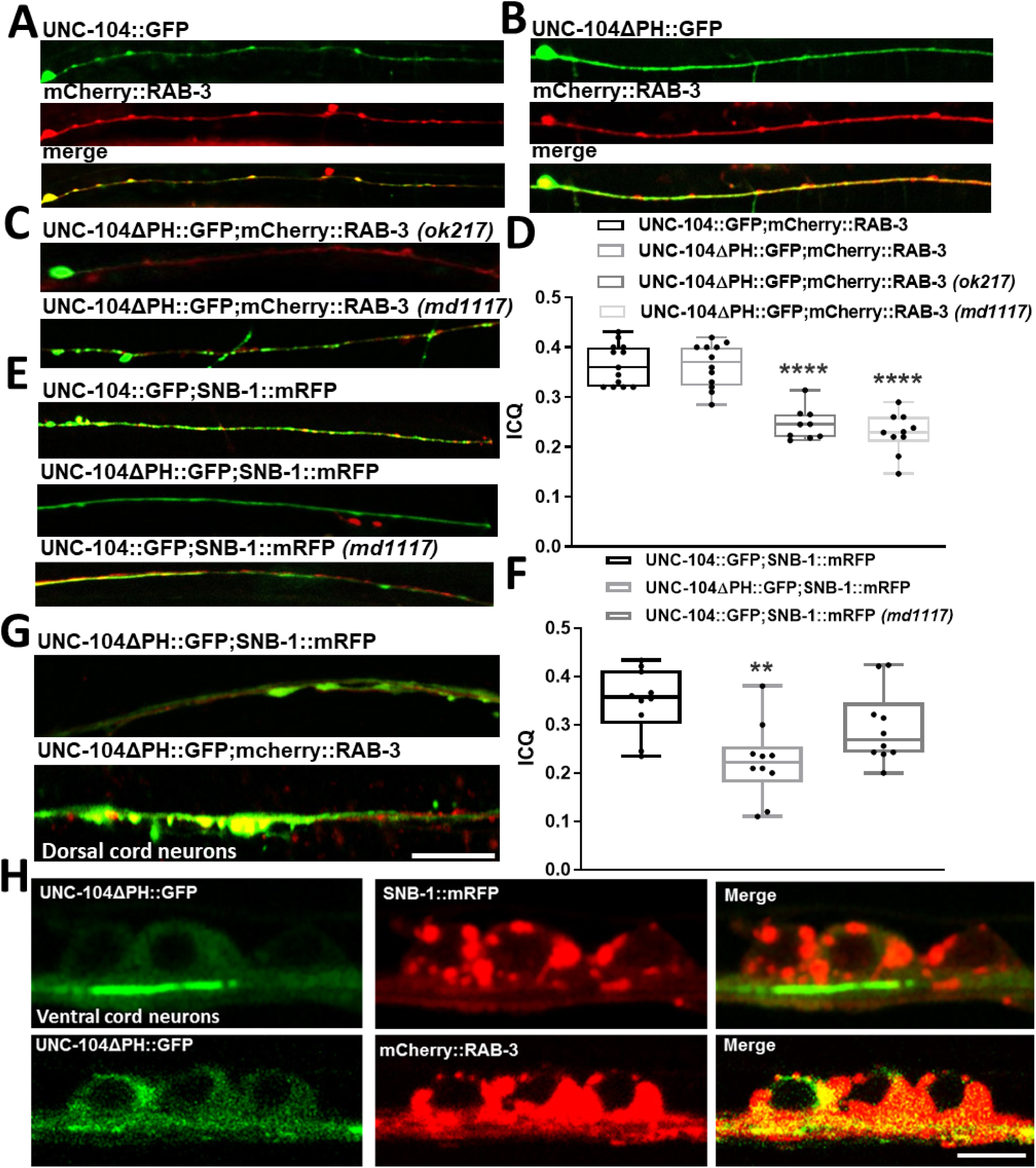
Colocalization between RAB-3/UNC-104 and SNB-1/UNC-104 depending on UNC-104’s PH domain and additional *unc-10* and *syd-2* mutations. (A+B) Colocalization between RAB-3 and (A) UNC-104 full length or (B) UNC-104 without its PH domain (UNC-104ΔPH). (C) Colocalization between UNC-104ΔPH and RAB-3 in *syd-2* or *unc-10* knockout worms. (D) Quantification of data from (A-C). For line scans along these sublateral neurons refer to Suppl. Fig. S4. (E) SNB-1 colocalization with full length UNC-104 (wt and *unc-10* mutant) or with UNC-104ΔPH. (F) Quantification of data from (E). (G-H) Colocalization of UNC-104ΔPH/SNB-1 and UNC-104ΔPH/RAB-3 in dorsal and ventral nerve cords. ANOVA with Dunnett’s multiple comparisons with **p<0.005, ***p<0.001, ****p<0.0001. Box and whisker graphs represent maximum value, upper quartile, median, lower quartile and minimum value. N = 20 worms for all experimental groups. Scale bar: 10 µm.

## DISCUSSION

To capture a synaptic transport vesicle at the pre-synapse and to position it in a way that it perfectly opposes the neurotransmitter receptors located at the post-synapse requires a complex protein machinery. Active zone proteins often inherit multiple binding partners and their multifaceted interaction sites and complex functions have been extensively studied. A key player in this protein complex is liprin-α(SYD-2) that encompasses binding sites for KIF1A(UNC-104), RIMS1(UNC-10), CAST, GIT, as well as the LAR(PTP-3) receptor. RIMS1(UNC-10) itself binds to RAB3A(RAB-3), liprin-α(SYD-2), Munc-13(UNC-13) and CAST (Zhen and Jin, 2004). Based on this knowledge, we propose a model in which RAB-3-containing vesicles (but not SNB-1-containing vesicles) would benefit from an additional linkage to support its binding to kinesin-3 UNC-104 (Fig. 7). This additional physical support would also strengthen the well-described motor-lipid linkage (Klopfenstein and Vale, 2004) which is considered to be rather weak and non-specific (Lemmon and Ferguson, 2001). How motors bind and recognize their cargo is vividly discussed in the literature (Brady and Morfini, 2017; Franker and Hoogenraad, 2013; Maday et al., 2014) and the dual UNC-10/SYD-2 complex may act as a novel receptor for UNC-104 to selectively transport RAB-3-containing vesicles. Our model is supported by findings that SYD-2/UNC-10, UNC-10/RAB-3 and SYD-2/RAB-3 all colocalize and are co-transported (Wu et al., 2013). Furthermore, liprin-α(SYD-2) also affects the transport of dense-core vesicles (Goodwin and Juo, 2013) and RIM1A(UNC-10) (known to be a dense-core vesicle protein) as well as RAB-3 are all mislocalized in *liprin-α/syd-2* mutants. Also, RIMB-1 and ELKS-1 localize independently of SYD-2 in dorsal nerve cords and this mechanism is affected by UNC-104 (Oh et al., 2021).

**Figure 7:**
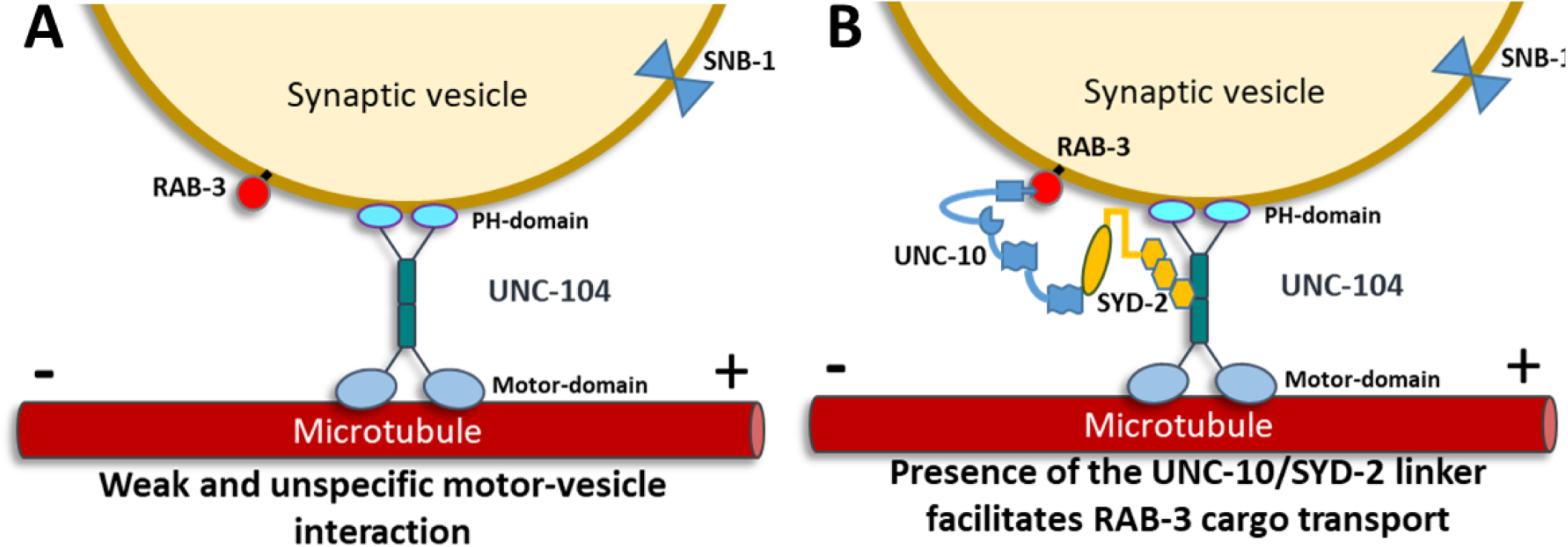
Model of an UNC-10/SYD-2 receptor in UNC-104-mediated RAB-3 cargo transport. (A) Current model of weak and unspecific UNC-104-vesicle interactions. (B) Presence of UNC-10/SYD-2 receptor stabilizes RAB-3-bound vesicles and promotes RAB-3 cargo transport.

Our analysis reveals a genetic linkage between unc-10 and syd-2 genes (Fig. 1A+B) as well as interdependencies between UNC-10 and SYD-2 to bind to UNC-104 (Fig. 1G+H). The physiological consequences of the UNC-10/SYD-2 linkage on UNC-104 neuronal particle distribution (Figs. 2+3), UNC-104 motility (Fig. 4), and UNC-104-mediated RAB-3 vesicle transport efficiencies (Figs. 5+6) are evident from this study. Indeed, it has been reported that UNC-10 acts to stabilize SYD-2 in active zone complexes and that UNC-10 binding to the C-terminus of SYD-2 facilitates the oligomerization of SYD-2 (Chia et al., 2013; Taru and Jin, 2011). The importance of oligomerized SYD-2 lies in its proposed function as a motor scaffolding protein that may bind to more than one UNC-104 at the same time (Chia et al., 2013; Wagner et al., 2009). Similarly, the functional relation between UNC-10/SYD-2 and RAB-3 is obvious from our experiments, e.g., SYD-2 binding to UNC-104 is significantly reduced in *rab-3*(*js49*); *unc-10 KD* double mutants (Fig. 1G+H). Moreover, the absence of *rab-3* negatively affects both UNC-104 neuronal distribution (Fig. 3C+E) as well as motor speeds, run lengths and pausing (Fig. 4); and these effects are significantly enhanced in the presence of *unc-10* mutations (Figs. 3 and 4).

UNC-104 requires ARL-8, a small GTPase to unlock its autoinhibitory state (Niwa et al., 2016). Because ARL-8 is usually associated with the cargo vesicle, it is possible that in the absence of *rab-3* not only the motor-cargo association is compromised, but also UNC-104 inactivation enhanced. Enhanced intramolecular folding of UNC-104 may then increasingly mask SYD-2 binding sites (such as SYD-2 SAM domain binding to UNC-104 FHA domain (Chia et al., 2013; Muniesh et al., 2020; Wagner et al., 2009)) consistent with our observations from co-immunoprecipitation assays (though not significant, there is an obvious tendency of decreased UNC-104 binding to SYD-2 in *rab-3* mutants, Fig. 1G+H). Critically, a study that dissected the direct interactions between RAB-3 and UNC-10 in *C. elegans* suggests epistatic regulation of these proteins (Gracheva et al., 2008). It has been also shown that in *syd-2* mutants UNC-10 and RAB-3 mislocalize to dendrites in DA neurons (Goodwin and Juo, 2013). Furthermore, loss of *syd-2*, *unc-10* and *rab-3*, all cause depletion of synaptic vesicles at the active zones (Stigloher et al., 2011). To interpret measured *rab-3* effects, we also need to consider the enhancing or compensating effects by other Rabs (that may explain the partial overlapping functions of RAB-3) such as RAB-27, a close homolog of RAB-3 regulated by AEX-3 (DENN/MADD) (Mahoney et al., 2006; Niwa et al., 2008). On the other hand, it has been shown that various synaptic vesicle precursors are transported in a RAB-independent fashion (Niwa et al., 2008) consistent with our observations that SNB-1 is transported independently of RAB-3 and vice versa (Fig. 5A+C). More critically, we have shown that SNB-1 transport is independent of UNC-10 (Figs. 5+6). Overall slower movement of SNB-1, as opposed to RAB-3 (Fig. 5A+C, wild type data), may also be explained by the ‘multiple motor model’ (Vershinin et al., 2007) in which multiple UNC-104 motors bind to a single RAB-3 vesicle to boost transport speeds.

Visible negative effects of *unc-10* mutations on UNC-104’s neuronal distributions (Fig. 3) and motor motility (Fig. 4) may lead to the notion that UNC-10 stabilizes the binding of SYD-2 to UNC-104 (Fig. 7). Loss of UNC-10 would then subsequently result in loss of SYD-2 binding leading to motor inactivation as well as reduced axonal clustering (Wagner et al., 2009; Wu et al., 2016). This notion is in accordance with observations that unc-10 knockdown even reduces observed SYD-2 overexpression effects (thus boosting motor speeds, Fig. 4A). Nevertheless, the concept of a dual UNC-10/SYD-2 linker to enhance binding specificity between UNC-104 and RAB-3-containing vesicles is only valid if deleting the motor’s PH domain would not (or only little) affect motor/RAB-3-vesicle interactions. Indeed, in worms carrying a mutation/deletion in the PH domain motor/RAB-3-vesicle interaction remains unaffected while this mutation significantly affects motor/SNB-1-vesicle interaction (Figs. 5+6, Suppl. Fig. S4). Besides the general assumption that the PH domain-binding to membranous phosphoinositides is weak (Lemmon and Ferguson, 2001), varying vesicular surface charges might modulate protein-lipid interactions (Richens et al., 2015). Therefore, although protein-lipid interaction seems to be imperative for UNC-104-vesicle interactions (Klopfenstein et al., 2002), both specificity and binding-strength of such interactions are generally low. Thus, the novel dual linker UNC-10/SYD-2 is critical to physically strengthen motor-vesicle interactions. Moreover, because UNC-104 undergoes dimerization upon membrane-cargo binding (Klopfenstein et al., 2002; Tomishige et al., 2002), such an additional linker complex may prolong the motor’s activated state.

## MATERIALS & METHODS

### *C. elegans* strains and plasmids

Worms were maintained at 20°C on NGM agar plates seeded with auxotroph *E. coli* strain OP50 (as a food source) according to standard methods (Brenner, 1974). Transgenic animals *Punc-104::unc-104::gfp*(*e1265*), *Punc-104::unc-104::mrfp*(*e1265*) and *Punc-104::unc-104::mrfp*(*ok217*) were described previously (Tien et al., 2011; Wagner et al., 2009). Strain *Punc-104::unc-104ΔPH::gfp*(*e1265*) is a kind gift from Dr. Dieter Klopfenstein (Georg-August-University, Göttingen, Germany) (see also (Klopfenstein and Vale, 2004)). Existing constructs *Punc-104::unc-104::gfp* and *Punc-104::unc-104::mRFP* were microinjected (using a standard protocol (Mello et al., 1991)) at a concentration of 100 ng/μl into NM1657 *unc-10*(*md1117*) and NM791 *rab-3*(*js49*) mutant worms to generate transgenic lines OIW 77 *unc-10*(*md1117*)*;nthEx77*[*Punc-104*::*unc-104::mrfp*], OIW 78 *unc-10*(*md1117*)*;nthEx78*[*Punc-104*::*unc-104::gfp*] and OIW 79 *rab-3*(*js49*)*;nthEx79*[*Punc-104*::*unc-104::mrfp*]. Note that for optimal operation of the Punc104 promoter (thus optimal UNC-104 expression) rather high injection dosages are needed (Tien et al., 2011; Wagner et al., 2009). Indeed, only unc-104 (full-length) plasmid concentrations above 70 ng/μl fully rescue the highly uncoordinated *unc-104*(*e1265*) phenotype (Klopfenstein and Vale, 2004) though neither ectopic expression nor any effects on worm behavior were seen.

All mutant strains were received from the Caenorhabditis Genetics Center (Minnesota, USA). For colocalization studies, existing plasmid *Punc-104::gfp::syd-2* (Wu et al., 2016) was co-injected along with *Punc-104::unc-104::mrfp* (each at a concentration of 75 ng/μl) into N2 wild type animals, *unc-10*(*md1117*) and *rab-3*(*js49*) mutants animals to obtain strains OIW 80 N2;*nthEx80*[*Punc-104::gfp::syd-2;Punc-104::unc-104::mrfp*], OIW 81 *unc-10* (*md1117*);*nthEx81*[*Punc-104::gfp::syd-2;Punc-104::unc-104::mrfp*] and OIW 82 *rab-3*(*js49*);*nthEx82*[*Punc-104::gfp::syd-2;Punc-104::unc-104::mrfp*]. Likewise, *Punc-104::gfp::syd-2* (75 ng/μl) was microinjected into *unc-10*(*md1117*)*;rab-3*(*js49*)[*Punc-104::unc-104::mrfp*] to obtain OIW 83 *unc-10*(*md1117*)*,rab-3*(*js49*); *nthEx83*[*Punc-104::gfp::syd-2;Punc-104::unc-104::mrfp*]. For BiFC (bimolecular fluorescence complementation) studies, the following strains were generated using existing plasmids *Punc-104::unc-104::VC155* and *Punc-104::VN173::syd-2* (Hsu et al., 2011): OIW 84 N2;*nthEx84*[*Punc-104::unc-104::VC155;Punc-104::VN173::syd-2*], OIW 85 *unc-*

*10*(*md1117*);*nthEx85*[*Punc-104::unc-104::VC155;Punc-104::VN173::syd-2*] and OIW 86 *rab-3*(*js49*)*;nthEx86*[*Punc-104::unc-104::VC155;Punc-104::VN173::syd-2*], respectively. For *rab-3* rescue strains, we designed a *Punc-104::rab-3::yfp* plasmid and injected it at 75 ng/μl into *rab-3*(*js49*)*;nthEx*[*Punc-104*::*unc-104::mrfp*] mutant to create OIW 87 *rab-3*(*js49*)*;nthEx87*[*Punc-104*::*unc-104::mrfp;Punc-104::rab-3::yfp*]. Likewise, *unc-10* rescue strains were generated by cloning *Punc-104::unc-10::gfp*and injecting it at 75 ng/μl into *unc-10*(*md1117*)*;nthEx*[*Punc-104*::*unc-104::mrfp*] mutant to create OIW 88 *unc-10*(*md1117*)*;nthEx88*[*Punc-104*::*unc-104::mrfp;Punc-104::unc-10::gfp*]. Similarity, *syd-2* rescue strains were obtained by co-injecting existing plasmid *Punc-104::gfp::syd-2* along with *Punc-104::unc-104::mrfp* (each at a concentration of 75 ng/μl) into *syd-2*(*ok217*) mutant worms to receive OIW 89 *syd-2*(*ok217*);*nthEx89*[*Punc-104::gfp::syd-2;Punc-104::unc-104::mrfp*]. *Punc-104::mcherry::rab-3* was designed by amplifying the rab-3 gene from cDNA libraries using primers ATGGCGGCTGGCG-GACAA (forward) and TTAGCAATTGCATTGCTGTT (reverse). In order to generate transgenic lines expressing cargo markers, we injected existing constructs *Punc-86::snb-1::mRFP* (Wagner et al., 2009) and *Punc-104::mcherry::rab-3* into N2 wild type animals, *unc-10*(*md1117*) and *syd-2*(*ok217*) mutants (at 100 ng/μl) to obtain strains OIW 90 N2;*nthEx90*[*Punc-86::snb-1::mRFP*], OIW 91 N2;*nthEx91*[*Punc-104::mcherry::rab-3*], OIW 92 *syd-2*(*ok217*)*;nthEx92*[*Punc-86::snb-1::mRFP*], OIW 93 *syd-2*(*ok217*)*;nthEx93*[*Punc-104::mcherry::rab-3*], OIW 94 *unc-10*(*md1117*)*;nthEx94*[*Punc-86::snb-1::mRFP*] and OIW 95 *unc-10*(*md1117*)*;nthEx95*[*Punc-104::mcherry::rab-3*]. We injected existing constructs *Punc-104::snb-1::mRFP* (80ng/µl) and *Punc-104::mcherry::rab-3* (80ng/µl) into *unc-104(e1265)*, respectively to obtain strains OIW 96 *e1265;nthEx96[Punc-104::snb-1::mRFP]* and OIW 97 *unc-104(e1265);nthEx97[Punc-104::mcherry::rab-3].* Also, the same construct *Punc-104::snb-1::mRFP* (80 ng/µl) and *Punc-104::mcherry::rab-3* (80 ng/µl) was injected into *unc-10(md1117)*, respectively, to obtain strains OIW 98 *unc-10(md1117);nthEx98[Punc-104::snb-1::mRFP]* and OIW 99 *unc-10(md1117);nthEx99[Punc-104:: mcherry::rab-3]*For UNC-104 and RAB-3 colocalization studies, plasmids *Punc-104::unc-104ΔPH::gfp* and *Punc-104::mcherry::rab-3* were coinjected into N2 wild type animals, *unc-10*(*md1117*) and *syd-2*(*ok217*) mutants to generate OIW 100 N2;*nthEx100*[*Punc-104::unc-104ΔPH::gfp;Punc-104::mcherry::rab-3*], OIW 101 *unc-10*(*md1117*);*nthEx101*[*Punc-104::unc-104ΔPH::gfp;Punc-104::mcherry::rab-3*] and OIW 102 *syd-2*(*ok217*)*;nthEx102*[*Punc-104::unc-104ΔPH::gfp;Punc-104::mcherry::rab-3*].For BiFC experiments *Punc-104::rab-3::VN173* and *Punc-104::snb-1::VC155* plasmids were generated by amplifying *rab-3* cDNA using primers GGCGCGCCATGAATAATCAACAGGC (forward) with an AscI site and ACCGGTGCAATTGCATTGCTGTTGAG (reverse) with an AgeI site and substituted a *map-1* insert in *Punc-104::map-1::VN173* (Wu et al., 2016) with *rab-3*. Similarly, *snb-1* genomic DNA was amplified using primers GGCGCGCCATGGACGCTCAAGGAGATGC (forward) with an AscI site and GGTAC-CTTTTCCTCCAGCCCATAAAACGATGA (reverse) with a KpnI site and substituted *lin-2* in *Punc-104::lin-2::VC155* (Wu et al., 2016) with *snb-1*. These plasmids were then co-injected (80 ng/μl) into N2 worms to generate OIW 103 N2;*nthEx103*[*Punc-104::rab-3::VN173;Punc-104::snb-1::VC155*]. Lastly, *unc-10*(*md1117*)*;rab-3*(*js49*) double mutant was generated by crossing *unc-10* (*md1117*)*;nthEx*[*Punc-104*::*unc-104::mrfp*] expressing male worms with *rab-3*(*js49*) hermaphrodites. Red fluorescent males from the F_1_ generation exhibiting coiler phenotypes were back-crossed into *rab-3*(*js49*) F_0_ hermaphrodites to obtain *unc-10*(*md1117*)*;rab-3*(*js49*) homozygous animals.

### Real-time PCR

To evaluate *syd-2* RNA levels in lysates from *unc-10* and *rab-3* mutant worms (Fig. 1A+B), we performed Real-Time PCR assays employing an ABI StepOnePlus Real-Time PCR machine in conjunction with the ABI Power SYBR green PCR master mix. Worms lysates are prepared from young adults based on a previously described protocol (Ward et al., 1988). RNA levels of *syd-2* were normalized to the *cdc-42* gene that acts as an internal control. The following primers were used: CGGAACAATACTCGACTTC (forward) and GCCACACGCTCCATT (reverse) that encompasses parts of the 2^nd^ and 4^th^ exon (200 bp) for *syd-2* and CTGCTGGACAGGAAGATTACG (forward) and TCGGACATTCTCGAATGAAG (reverse) for *cdc-42*.

### RNAi feeding assay

For RNA interference experiments, we employed the RNAi feeding method (Kamath et al., 2001) in which worms are fed by bacteria producing the desired dsRNA. We obtained feeding clones UNC-10(X-3F06), SYD-2(X-5K09) and SNB-1 (V-4J14) from the Julie Ahringer’s *C. elegans* RNAi feeding library (Source BioScience LifeSciences, USA; a kind gift of the *C. elegans* Core Facility, Taiwan) which we sequenced to determine their correctness. NGM plates containing ampicillin (25 μg/ml) and 1 mM IPTG were inoculated with the respective HT115 *E. coli* strain carrying the appropriate gene insert (flanked by T7 promoters) and grown overnight. 15-20 worms were transferred to the respective RNAi feeding plate and incubated at 20°C. Worms were then transferred to new RNAi feeding plates every 24 hours and the F1 progeny was used for analysis after day 5.

### Western blotting and co-immunoprecipitation assays

To extract proteins from worms, we prepared lysates from young adults based on a previously described protocol (Ward et al., 1988). To perform Western blots from whole worm lysates, 100 µg sample protein was used and membrane blocking was carried out using milk powder for 1-1.5 h at RT. Primary goat (N-terminal) anti-SYD-2 antibody (cL-19 #sc-15655, Santa Cruz) was incubated at 4°C for 14 h at 1/500 dilutions. Secondary anti-goat antibody was incubated at RT for 2 h with 1/1000 dilutions. Anti-β-actin antibody (sc-47778, Santa Cruz) was used as a loading control. For Co-IPs, we use the PureProteome Protein G Magnetic Bead System (EMD Millipore Corp.). 1 mg protein extract was used for immunoprecipitation and incubated with protein G beads and 4 µg anti-UNC-104 antibody (rabbit, polyclonal, commissioned from GeneTex based on an epitope designed by us). Western blotting (Fig. 1E+F) was performed using a 1:500 dilution of the anti-SYD-2 antibody (primary antibodies dilutions based on the manufacturer’s suggestion). Band density analysis was carried out using NIH ImageJ 1.50 software based on a published protocol (Schneider et al., 2012).

### Worm imaging and motility analysis

For microscopic observations, worms were immobilized on 2% agarose-coated cover slides, and no anesthetics (e.g., levamisole) were used (reported to affect motor motility (Kumar et al., 2010)). A Zeiss LSM780 confocal laser scanning microscope was employed for imaging worms as shown in Figures 2 and 6. For motility analysis (Fig. 4) and further imaging (Figs. 3 and 5), we employed an Olympus IX81 microscope with a DSU Nipkow spinning disk unit connected to an Andor iXon DV897 EMCCD camera for high-speed and long-duration time-lapse imaging (4-5 frames per second). To convert recorded time-lapse sequences into kymographs, we use imaging analysis software NIH ImageJ. The ‘straighten plugin’ was employed to straighten curved axons, and after drawing a line over the axon, the plugin ‘reslice stack function’ was executed to generate kymographs. In kymographs, static particles appear as vertical lines, whereas the slope of a moving particle corresponds to the velocity (speed) of the particle (Suppl. Fig. S1D). A pause is defined if motors move less than 0.07 μm/s, and each calculated velocity event does not contain any pauses. In average, we analyzed ≈600 moving events per experiment from 15 to 25 individual worms. A moving event is defined as a single motility occurrence typically right after a pause or a reversal, and an event ends when the motor again pauses or reverses (Suppl. Fig. S1D). Motor cluster analysis (Fig. 3B+E) was carried out using ImageJ’s ‘area’ tool and the ‘particle analyze’ plugin. To measure travel distances (Figs. 3D and 5B+D) of particles in (straightened) neurons, we used ImageJ’s ‘line tool’ to draw a line from the proximal axon hillock to the identified distal particle. Intensity Correlation Quotient (ICQ) (Figs. 2B and 6D+F) was determined by selecting the region of interest using the ‘polygonal selection tool’ in ImageJ. After background fluorescent intensity subtraction (ImageJ ‘subtract background’ function), the plugin ‘intensity correlation analysis’ was used to generate ICQ values. ICQ values range from −0.5 to 0.5, and values close to 0.5 stand for interdependent expression of two fluorophores while values close to −0.5 stand for segregated expression, and values around 0 imply random expression. For fluorescence intensity quantification (Fig. 2D), the following formula was used: ‘Integrated density of selected region – (area of selected region * mean fluorescence of the background)’ (Li et al., 2004). Parameters such as mean grey value of background, area of selected region and integrated density of selected region were determined using ImageJ’s ‘region selection’ tools. Line scans (Suppl. Fig. S4) were generated using ImageJ ‘plot profile’ function (after straightening the neurite using ImageJ’s ‘straighten’ tool).

## ACKNOWLEDGEMENTS

We thank the *C. elegans* Core Facility (CECF) Taiwan (funded by the Ministry of Science and Technology, MOST) for providing microinjection setups and worm observation systems. We acknowledge MOST grant NSC 100-2311-B-007-004 to OIW and support from the Brain Research Center of National Tsing Hua University under the Higher Education Sprout Project funded by the Ministry of Science and Technology and Ministry of Education in Taiwan.

## SUPPLEMENTARY MATERIAL

### SUPPLEMENTARY FIGURE

**Suppl. Figure S1:**
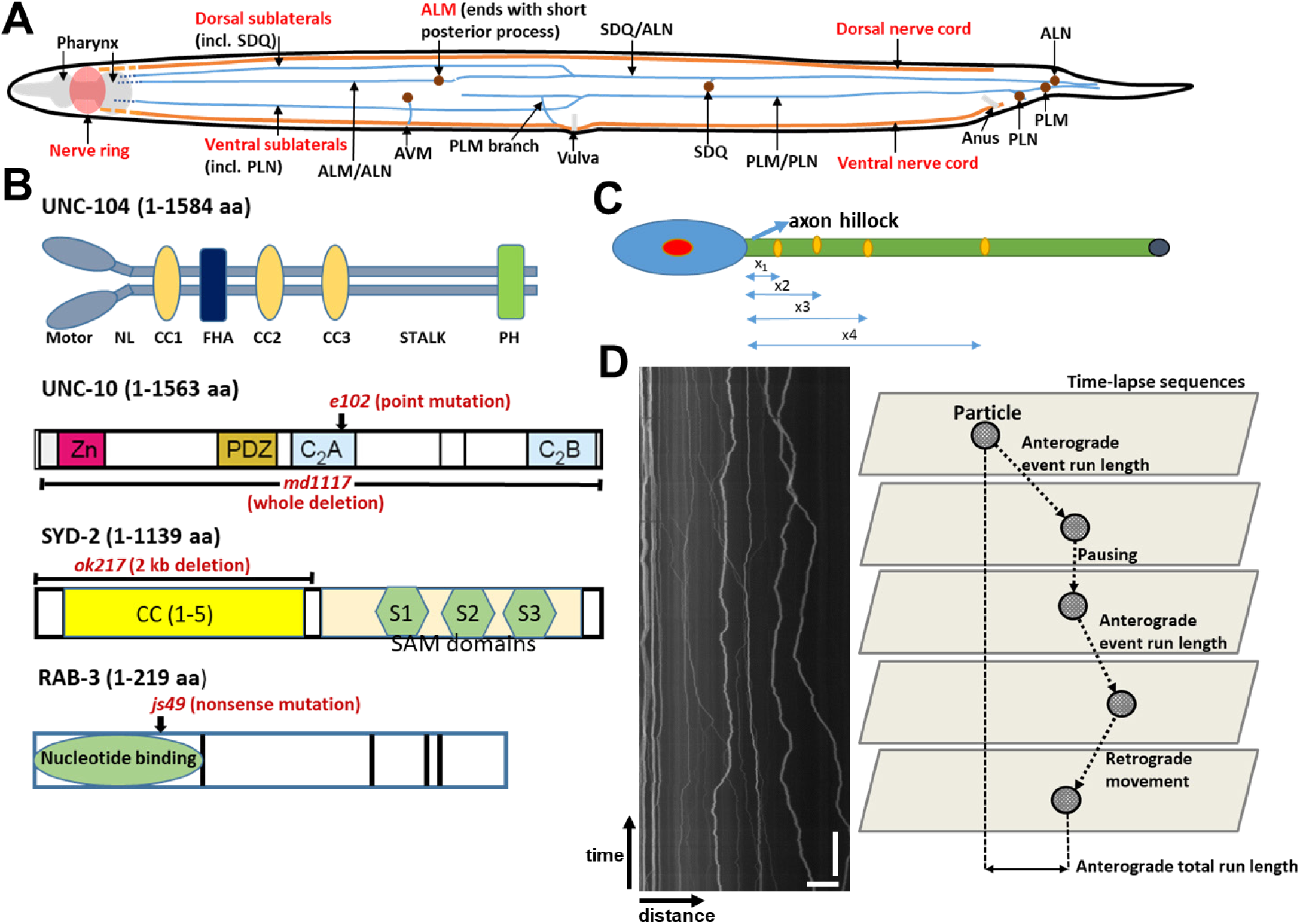
Diagrammatic schemes of used neurons, alleles and kymograph method. (A) Simplified scheme of the nervous system in *C. elegans* (ALM, sublaterals and nerve ring labelled in red). (B) Protein domain representation of UNC-104, UNC-10, SYD-2 and RAB-3. Mutant alleles marked in red. (C) Schematic diagram of a neuron with blue double-sided arrows indicating the distances travelled from the axon hillock to farther distal regions. (D) Kymograph and diagrammatic scheme of various motility parameters. Scale bars: vertical = 30 s, horizontal = 10 µm.

**Suppl. Figure S2:**
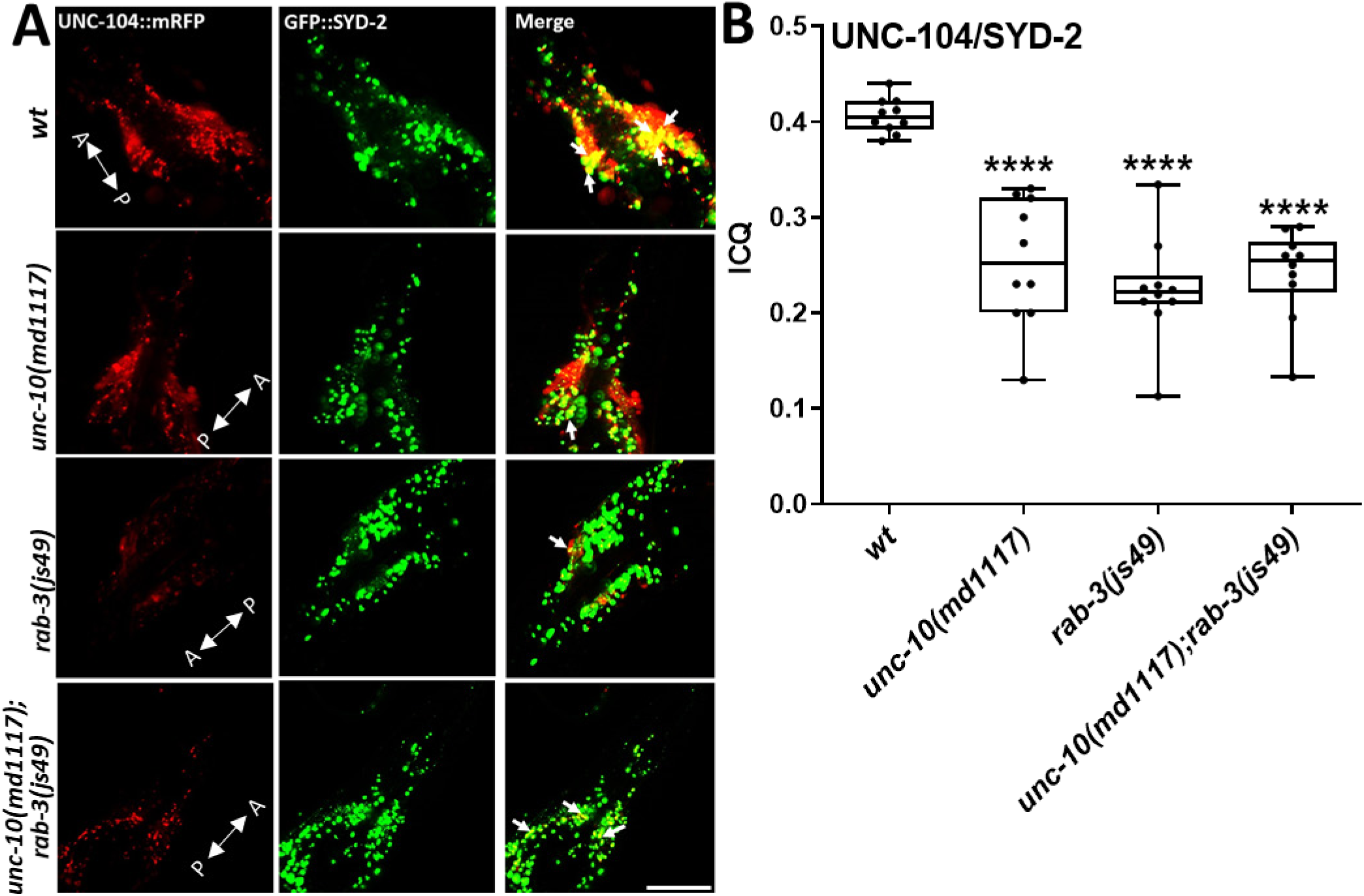
Colocalization between UNC-104 and SYD-2 in the presence or absence of *unc-10* or *rab-3* in nerve rings. (A) Representative images of UNC-104 and SYD-2 in *unc-10, rab-3* and *unc-10(md1117);rab-3(js49).* A = anterior, P = posterior. White arrowheads indicate particular colocalization events in nerve rings. (B) Intensity correlation coefficients from images shown in (A). N = 20 worms. One-way ANOVA with Dunnett’s test for multiple comparisons with *p<0.05, **p<0.005, ***p< 0.001, **** p<0.0001. Box and whisker graphs represent maximum value, upper quartile, median, lower quartile and minimum value. Scale bar: 10 µm.

**Suppl. Figure S3:**
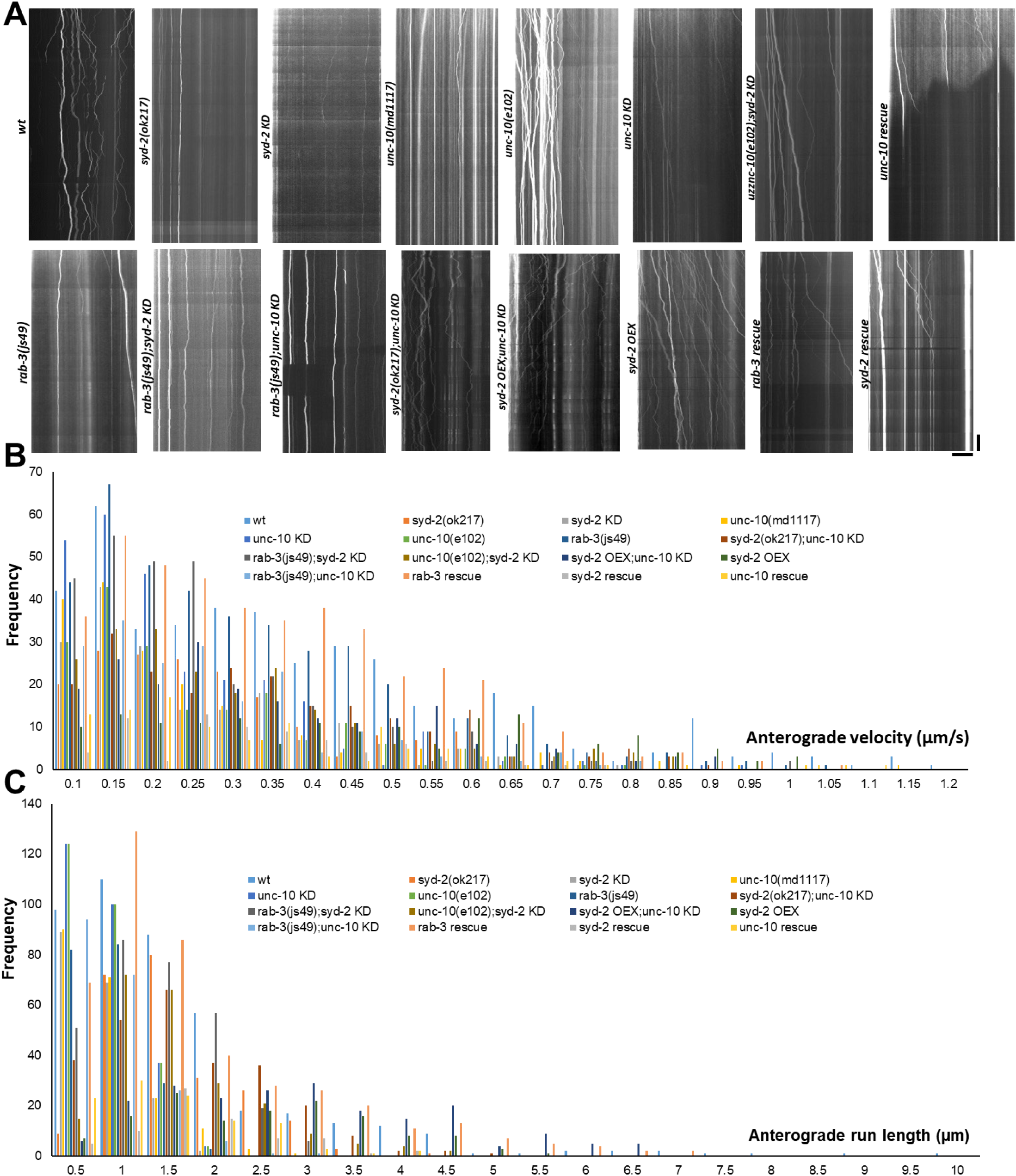
Representative kymographs as well as velocity and run lengths frequency distributions. (A) Representative kymographs from worms expressing UNC-104::mRFP in various strains. (B+C) Anterograde velocity and run length frequency distribution from data shown in Fig 4. A+B. Vertical scale bar: 25 s. Horizontal scale bar: 10 µm.

**Suppl. Figure S4:**
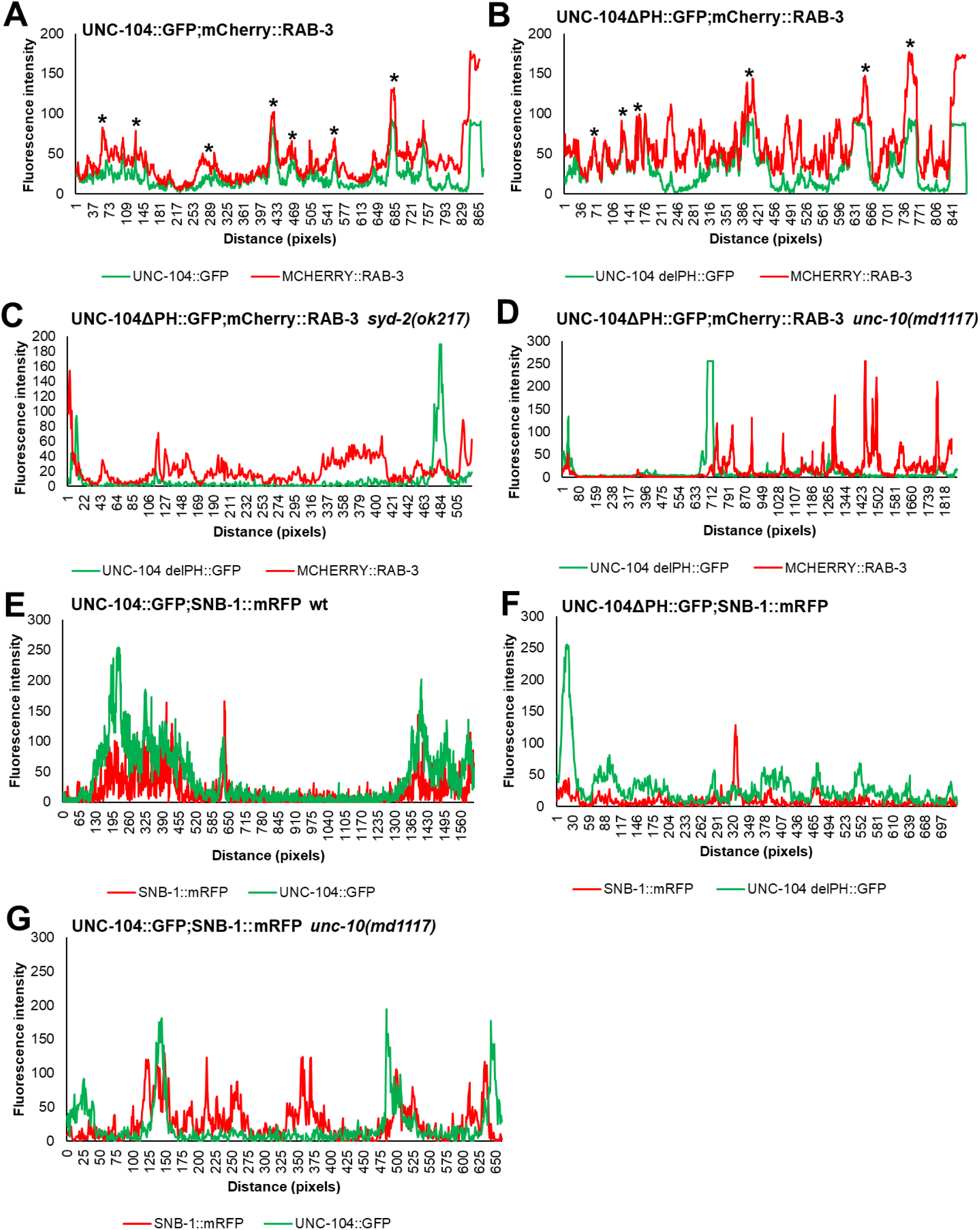
Line scans on sublateral neurons in worms expressing RAB-3 and UNC-104 (with and without PH domain) in wild type and various mutants. (A) Line scans along sublaterals of worms expressing UNC-104::GFP (green line) and mCherry::RAB-3 (red line). Note the overlapping (higher) peaks representing occurrences of colocalization (marked by wildcards). (B) Even when UNC-104’s PH domain is deleted UNC-104/RAB-3 peaks still overlap. (C+D) However, if additionally (C) *syd-2* or (D) *unc-10* is mutated UNC-104/RAB-3 peaks are largely separated. (E) Line scans along sublaterals of worms expressing UNC-104::GFP (green line) and SNB-1::mRFP (red line). (F) When UNC-104’s PH domain is deleted fluorescence intensity peaks become more separated and detached. (G) When additionally mutating *unc-10* the effect remains comparable.

### SUPPLEMENTARY TABLE

**Supplementary Table S1:** Accompanying table with details to Figure 3B.

| UNC-104::mRFP | wt | *syd-2(ok217)* | *unc-10(md1117)* | *rab-3(js49)* |
| --- | --- | --- | --- | --- |
| Average cluster size ($\mu\text{m}^2$) | 0.72±0.64 | 0.474±0.38 | 0.574±0.48 | 0.655±0.56 |
| Total neurite length (mm) | 1.95 | 2.44 | 1.44 | 1.28 |
| Cluster per 100 $\mu\text{m}$ | 19.55 | 15.5 | 22.7 | 25.4 |
| # cluster | 382 | 380 | 330 | 326 |

**Supplementary Table S2:** Accompanying table with details to Figure 3D+E.

| | Average distance ($\mu\text{m}$) | Cluster size ($\mu\text{m}^2$) |
| --- | --- | --- |
| UNC-104::mRFP | Mean±SEM | Mean±SEM |
| wt | 40.77±1.510 | 4.458±1.253 |
| *syd-2(ok217)* | 34.26±1.563 | 1.445±0.189 |
| *unc-10(md1117)* | 35.60±1.602 | 0.883±0.119 |
| *rab-3(js49)* | 39.63±1.432 | 1.021±0.413 |
| *rab-3(js49);syd-2 KD* | 54.41±2.866 | 0.898±0.177 |
| *rab-3(js49);unc-10(md1117)* | 9.809±1.255 | 0.976±0.420 |
| *syd-2(ok217);unc-10 KD* | 29.30±2.442 | 1.702±0.502 |
| *syd-2 rescue* | 35.99±2.472 | 2.522±0.383 |
| *unc-10 rescue* | 36.07±2.346 | 2.304±0.486 |
| *rab-3 rescue* | 40.01±3.316 | 2.599±0.458 |

**Supplementary Table S3:** Accompanying table with details to Figure 4A-D.

| UNC-104::mRFP | Anterograde velocity ($\mu\text{m/s}$) | | Anterograde run-length ($\mu\text{m}$) | | Anterograde total run length ($\mu\text{m}$) | | Pausing 10 $\mu\text{m}$ | |
| --- | --- | --- | --- | --- | --- | --- | --- | --- |
|  | Mean | N | Mean | N | Mean | N | Mean | N |
| wt | 0.3558 | 424 | 1.414 | 424 | 11.14 | 51 | 3.092 | 30 |
| *syd-2(ok217)* | 0.2526 | 178 | 1.375 | 236 | 12.53 | 20 | 5.854 | 22 |
| *syd-2 KD* | 0.2325 | 183 | 0.6156 | 183 | 4.172 | 27 | 11.78 | 26 |
| *unc-10(md1117)* | 0.2475 | 198 | 0.6791 | 199 | 5.631 | 24 | 8.220 | 22 |
| *unc-10(e102)* | 0.3339 | 217 | 1.627 | 223 | 13.14 | 14 | 4.244 | 17 |
| *unc-10 KD* | 0.2150 | 264 | 0.6401 | 265 | 6.524 | 26 | 3.680 | 66 |
| *unc-10(e102);syd-2 KD* | 0.2849 | 217 | 1.364 | 223 | 11.27 | 27 | 3.686 | 22 |
| *rab-3(js49)* | 0.2934 | 400 | 0.6509 | 198 | 5.603 | 23 | 9.310 | 24 |
| *rab-3(js49);syd-2 KD* | 0.2409 | 297 | 1.119 | 296 | 10.88 | 22 | 4.026 | 23 |
| *rab-3(js49);unc-10 KD* | 0.2476 | 192 | 0.6443 | 200 | 4.444 | 29 | 8.753 | 30 |
| *syd-2(ok217);unc-10 KD* | 0.3214 | 223 | 1.469 | 264 | 13.92 | 16 | 3.575 | 29 |
| *syd-2 OEX;unc-10 KD* | 0.3214 | 207 | 2.664 | 210 | 12.69 | 15 | 1.170 | 28 |
| *syd-2 OEX* | 0.4380 | 155 | 2.244 | 138 | 18.78 | 11 | 0.8253 | 23 |
| *rab-3 rescue* | 0.3234 | 431 | 1.573 | 442 | 12.26 | 43 | 4.339 | 42 |
| *syd-2 rescue* | 0.2996 | 74 | 1.593 | 74 | 14.31 | 4 | 2.284 | 6 |
| *unc-10 rescue* | 0.2940 | 98 | 1.240 | 110 | 11.06 | 9 | 4.939 | 10 |

**Supplementary Table S4:** Accompanying table with details to Figure 5A+C.

| SNB-1::mRFP | Anterograde velocity ($\mu\text{m/s}$) | |
| --- | --- | --- |
|  | Mean | N |
| wt | 0.317 | 158 |
| *syd-2(ok217)* | 0.230 | 159 |
| *unc-10(md1117)* | 0.315 | 203 |
| *unc-10(md1117);syd-2 KD* | 0.270 | 72 |
| *rab-3 KD* | 0.310 | 149 |
| *unc-104(e1265)* | 0.220 | 199 |
| **mCherry::RAB-3** |  |  |
| wt | 0.351 | 174 |
| *syd-2(ok217)* | 0.226 | 158 |
| *unc-10(md1117)* | 0.275 | 182 |
| *unc-10(md1117);syd-2 KD* | 0.233 | 74 |
| *snb-1 KD* | 0.367 | 184 |
| *unc-104(e1265)* | 0.253 | 189 |

**Supplementary Table S5:**
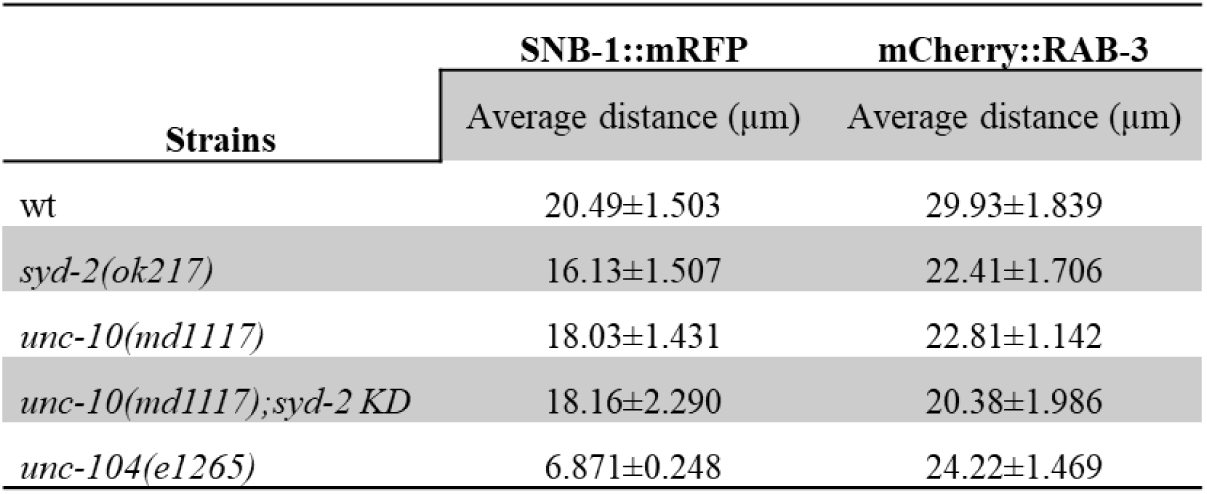
Accompanying table with details to Figure 5B+D.

